# Comprehensive Characterisation of Fetal and Mature Retinal Cell Identity to Assess the Fidelity of Retinal Organoids

**DOI:** 10.1101/2022.06.13.495996

**Authors:** Hani Jieun Kim, Michelle O’Hara-Wright, Daniel Kim, To Ha Loi, Benjamin Y. Lim, Robyn V. Jamieson, Anai Gonzalez-Cordero, Pengyi Yang

## Abstract

Characterizing cell identity in complex tissues such as the human retina is essential for studying its development and disease. While retinal organoids derived from pluripotent stem cells have been widely used to model development and disease of the human retina, there is a lack of studies that have systematically evaluated molecular and cellular fidelity of the organoids derived from various culture protocols in recapitulating their *in vivo* counterpart. To this end, we performed an extensive meta-atlas characterisation of cellular identities of the human eye, covering a wide range of developmental stages. The resulting map uncovered previously unknown biomarkers of major retinal cell types and those associated with cell-type specific maturation. Using our retinal cell identity map from the fetal and adult tissues, we systematically assessed the fidelity of the retinal organoids to mimic the human eye, enabling us to comprehensively benchmark the current protocols for retinal organoid generation.

## Introduction

The human retina is a complex tissue and comprises various cell types that function together to convert light into biological signals. To understand the development and disease of the human eye requires the characterization of molecular and cellular programs that define the identity of cells in the human retina. While the access to the human retina tissues is limited, human retinal organoids derived from pluripotent stem cells (PSCs) which aim to closely recapitulate the human eye offer unprecedented opportunities to investigate early retinal development and disease. Furthermore, an increasing number of studies are harnessing the potential of retinal organoids for therapeutic applications such as cell transplantation and as a model to evaluate the efficacy of treatment strategies (Chahine Karam et al., 2022; Ribeiro et al., 2021; West et al., 2022). Early studies have used human embryonic stem cell (hESC)-derived embryoid bodies followed by two-dimensional (2D) culturing to generate retinal precursors, which were then isolated for directed and undirected differentiation into ganglion and amacrine cells, and to a lesser extent photoreceptor precursor cells (Lamba et al., 2006). Whilst the efficient production of the retinal progenitors under 2D conditions enabled useful initial applications in cell therapy studies (Meyer et al., 2009; Osakada et al., 2008), these systems lack the capacity to recapitulate the three-dimensional (3D) features of the native retinal cells *in vivo*. This has led the field to develop advanced 3D *in vitro* structures that can recapitulate the physiological, morphological, and spatiotemporal patterns of the developing retina. These were first developed using mouse embryonic stem cells (Eiraku et al., 2011) and later translated to hESCs to generate the so-called human retinal organoids (Lamba et al., 2006; West et al., 2012).

Since then, several advances in the retinal organoid field have led to the development of the state-of-the-art protocols that allow efficient and rapid formation of retinal organoids consisting all the retinal cell types: ganglion, amacrine, bipolar, horizontal, Müller glial, and photoreceptor cells (Afanasyeva et al., 2021). These organoids have been shown to generate mature features such as ribbon synapses (Artero Castro et al., 2019) and outer segments with physiological response to light stimuli (Wahlin et al., 2017; Zhong et al., 2014), showing remarkable functional similarity to the eye (Gonzalez-Cordero et al., 2017). A mixture of 2D and 3D protocols that do not require the addition of small molecules allow generation of mature and light-sensitive photoreceptors with rudimentary outer segments (Zhong et al., 2014). Other protocols that involve stepwise a 2D-to-3D culture enable the formation of the embryoid body to be bypassed (Reichman et al., 2014). Other protocols have incorporated differentiation factors such as serum, RA, taurine, and supplements N2 and B27 (Gonzalez-Cordero et al., 2017) and antioxidants and lipids (West et al., 2022) that have significantly improved the generation of photoreceptor outer segments. Most of these protocols share common media components, such as BMP4 and IGF-1, but differ in their timing in the switch from 2D to 3D culture and/or the addition of certain molecules.

Increasing number of studies have begun profiling the human organoids derived from these protocols at single-cell resolution to investigate retinal development and disease (O’Hara-Wright and Gonzalez-Cordero, 2020). These studies have provided unprecedented opportunity to investigate the heterogeneity of the retinal cell types and uncovered several new insights into retinal development, such as the discovery of a potentially novel regulator of cone fate (Kallman et al., 2020), a population of post-mitotic transitional cells (Sridhar et al., 2020), and the convergence of retinal organoid transcriptomes towards peripheral retinal cell types (Cowan et al., 2020). Whilst these advancements have enhanced our understanding of retinal biology, the growing single-cell resource of the human retina and retinal organoids has yet been probed to systematically benchmark the state-of-the-art protocols for their capacity to produce organoids fidel to their *in vivo* counterpart.

Here, we performed an extensive curation of scRNA-seq datasets from the human retinal tissue and organoids derived from a variety of differentiation protocols, generated a comprehensive map of retinal cellular identities of the mature and fetal eye, and benchmarked the fidelity of the human retinal organoid models to faithfully recapitulate the human eye. The extensive meta-atlas characterisation of the retinal cellular identities enabled the discovery of an array of previously unknown marker genes of retinal cell types and those associated with cell-type-specific retinogenesis. Moreover, these cellular identities resolved by age were used to systematically benchmark the current protocols for their capacity to generate cell types that closely emulate their *in vivo* counterparts in terms of cell identity, cell-type proportion, and coverage. Finally, we developed a user-friendly application called Eikon (https://shiny.maths.usyd.edu.au/Eikon/) that facilitates users to assess the fidelity of their retinal organoids.

## Results

### Generating a cell identity map of the human retina

Single-cell transcriptome profiling has been applied to resolve the cellular identities in the retinal tissue (**Figure 1A**). To create a cell identity map of the human retina, we began by compiling a collection of scRNA-seq datasets generated from the human retinal tissue in “mature” samples including those from the postnatal stage and the adult retina (**Figure 1B**) (Cowan et al., 2020; Lu et al., 2020; Lukowski et al., 2019). These datasets provide a large number of cells (ranging from 7,382 to 32,375) and good coverage of genes and reads per cell with a median gene and read number of >1,000 in most datasets and samples from Cowan et al. (Cowan et al., 2020) showing the highest quantification rates per cell (**Figure S1A**).

**Figure 1.**
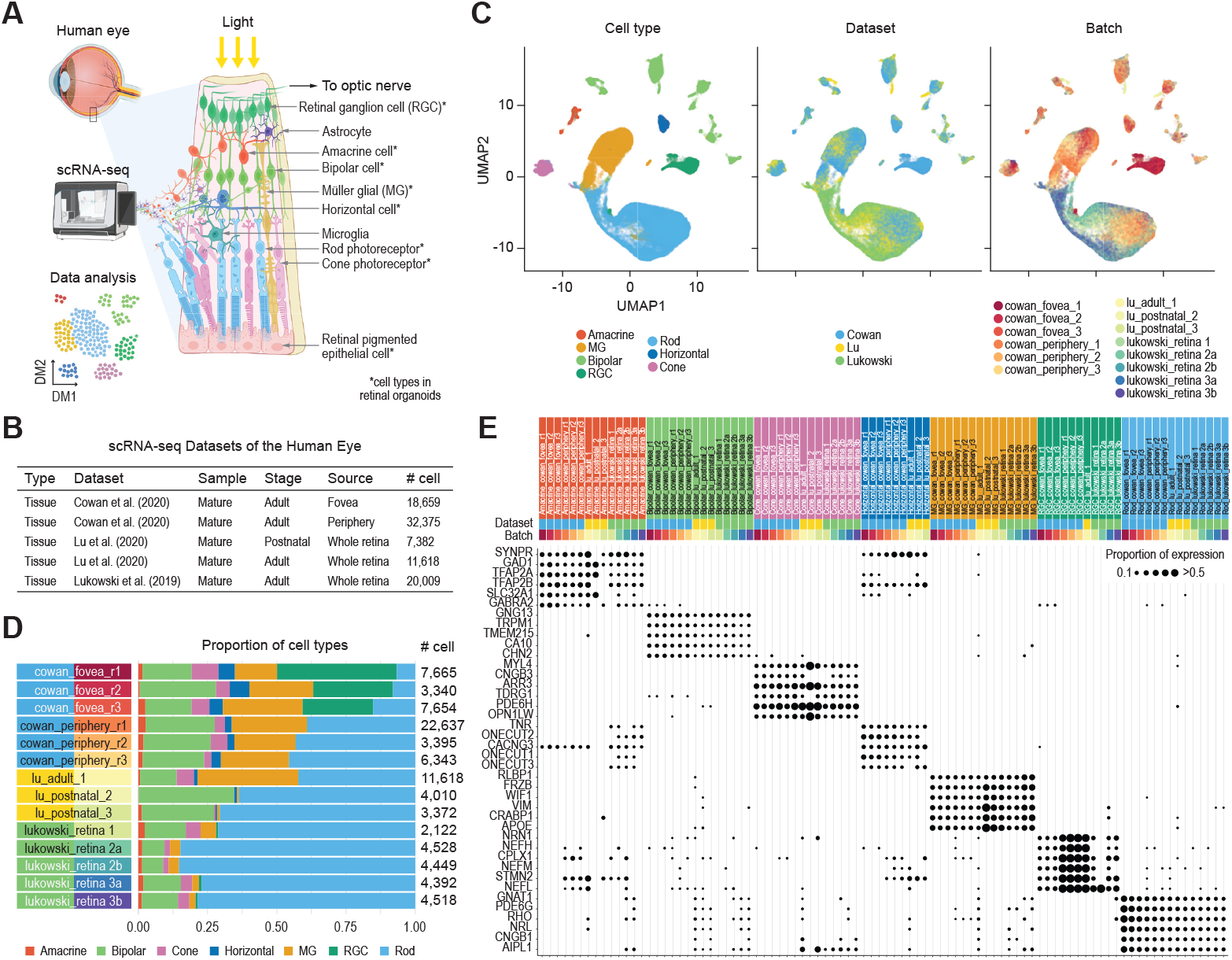
Curation of scRNA-seq datasets generated from the mature retinal tissue. **(A)** Schematic of single-cell transcriptomic profiling by scRNA-seq and the major cell types of the human retina. **(B)** Summary of scRNA-seq datasets collected from the mature retinal tissues. **(C)** UMAP representation of the transcriptomes of single cells. Cells are coloured by their type (left), dataset of origin (middle), and batch in each dataset (right). **(D)** Proportion of cell types (color coded) and total number of cells in each batch and dataset. **(E)** Expression pattern of known retinal cell type marker genes across datasets and batches.

We visualized the integrated datasets by cell type, dataset, and batch within each dataset using UMAP (**Figure 1C**). Next, we analyzed the number of cells in each cell type and their proportions in each dataset and batch (**Figure 1D and S1B**). We observed that, whilst most of the major cell types are identified in the datasets and the proportion of cell types are largely consistent across batches within each dataset, the proportion of some cell types showed large variability across datasets. The variability in proportion (such as those of the retinal ganglion and Müller glial cells) may be due to differences in the stage of tissue collection (e.g., adult versus postnatal stage) and/or to differences in the collection source (e.g., whole retina versus a specific location of the retina [fovea versus periphery]), as well as tissue processing time post-mortem. Consistent with the anatomical location of the fovea in the retina lacking the rod cells, the three samples harvested from the fovea demonstrated the lowest proportion of rods. Evaluating known retinal cell type gene markers showed that their expression is highly cell type specific (**Figure 1E**), confirming the accuracy in cell type annotation in the public datasets. Together, this large resource of scRNA-seq datasets profiling the human eye form the basis for the curation and characterization of the human retina cell types and for assessing the retinal cell identity and the fidelity of human retinal organoids to mimic the *in vivo* identities.

### Deriving robust cell identity scores of genes for retina cell types

To resolve genes that underlie retinal cellular identity, we computed a cell-type-specific cell identity score for each gene by dataset and batch using Cepo, a computational method for detecting cell identity genes (Kim et al., 2021). The clustering of samples from across datasets and batches using Pearson’s correlation of Cepo-derived gene statistics show strong grouping by cell type irrespective of the origin of dataset and batch (**Figure 2A**).

**Figure 2.**
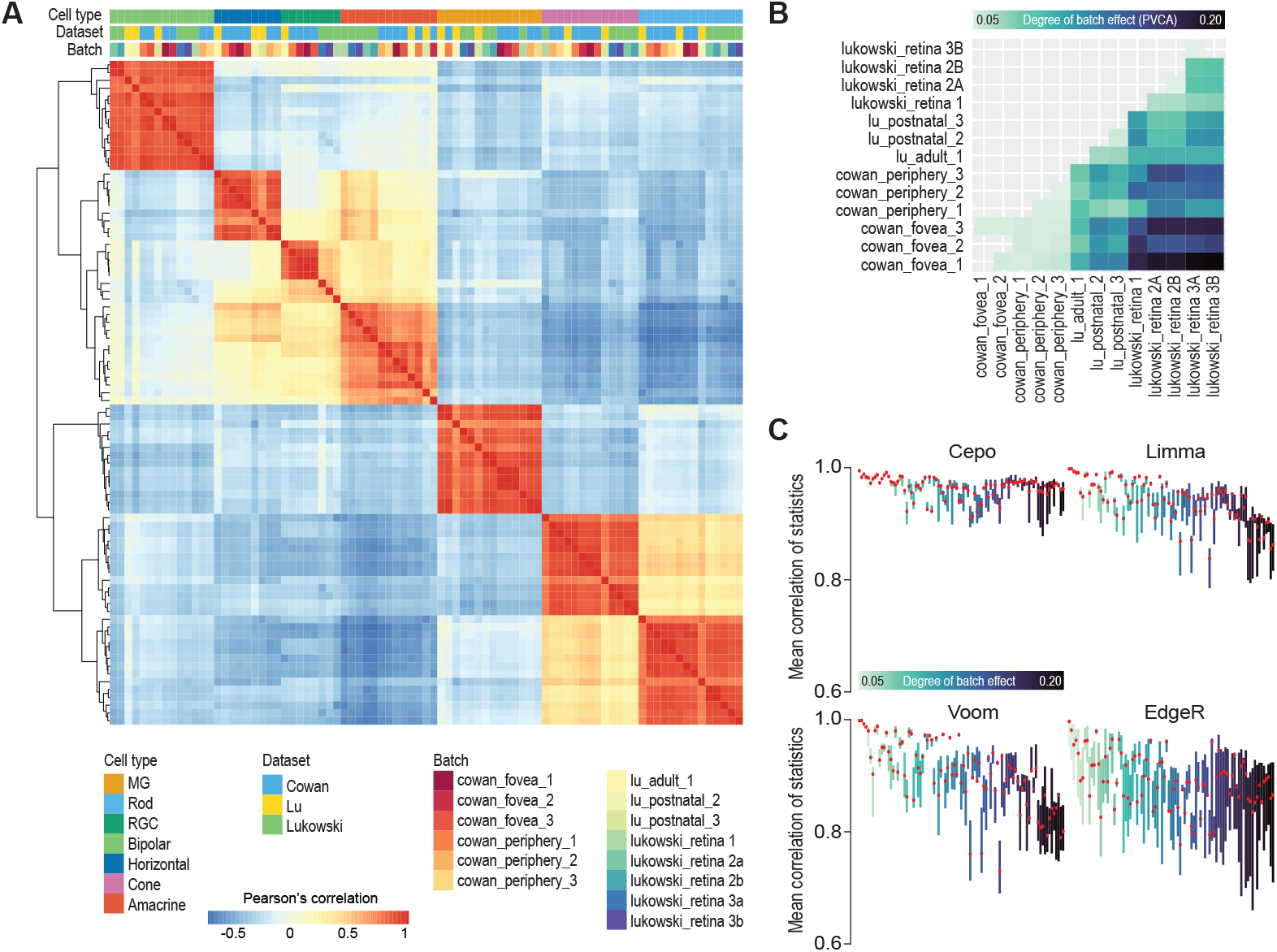
Comparative analysis of cell-type-specific gene statistics across datasets and batches. **(A)** Correlation heatmap of cell identity gene statistics generated from Cepo (Kim et al., 2021) for each cell type across datasets and batches. The heatmap is hierarchically clustered by the similarity of correlation profiles. **(B)** Pairwise assessment of batch effect using principal variance component analysis (PVCA). The proportion of variance contributed by batch in each pair of datasets is visualized. A darker color denotes stronger batch effect. **(C)** Boxplot of mean correlation of gene statistics from all retinal cell types for each pair of datasets illustrated in (B) using Cepo, Limma, Voom, and EdgeR statistics.

To systematically quantify the influence of the batch source on Cepo-derived cell identity gene statistics, we conducted two assessments. First, we applied hierarchical, Louvian, and *k*-means clustering on the samples and compared their concordance with respect to three sources of variation – cell type, dataset, or batch – using adjusted Rand index (ARI) and normalized mutual information (NMI). We found that the Cepo-derived cell identity gene statistics enabled accurate clustering of samples by their cell type label whereas both dataset and batch source had minimal influence on the clustering (**Figure S2A**). These findings were consistent across a varying number of genes (from top 10 to 100 genes per cell type) selected for the clustering (**Figure S2A**). These results demonstrate the high reproducibility of the cell-type identity statistics calculated using Cepo for genes across the retina scRNA-seq datasets and batches.

Next, we performed principal variance component analysis (PVCA) (Li et al., 2009) on all pairs of batches across datasets to quantify the degree of variance contributed by batch source (**Figure 2B**). Computing the degree of batch effect present in all pairs of batches generates a set of batch pairs ranging from those that exhibit low batch effect to those that exhibit high batch effect. As expected, batch pairs both originating from the same dataset demonstrate lower batch effect, whilst those originating from different datasets demonstrate higher batch effect. The batch effect was greatest between the batches from the Cowan dataset and those from the Lukowski dataset. We then computed the concordance of Cepo-derived gene statistics between the dataset pairs for each of all cell types and then evaluated these statistics against the degree of batch effect present in the data (**Figure 2C and S2B**). We found that the concordance in Cepo-derived gene statistics between the same cell types was retained across increasing batch effects (**Figure S2B**, first column). We compared the robustness of other measures of cell-type-specific gene statistics (Limma, Voom, and EdgeR) against batch effects. In contrast to the findings from Cepo statistics, we found that there was a gradual loss in concordance with increasing presence of batch effect across most cell types, with the most pronounced decrease in the statistics for the amacrine cells (**Figure S2B**, second to fourth columns). Collectively, the results from the batch assessment strongly support the high reproducibility of the Cepo-derived cell identity gene statistics across the retina scRNA-seq datasets, highlighting their robustness against batch effect.

### The retinal cell identity map uncovers novel cell identity genes

To uncover potentially new markers of cell types, we performed a systematic query search on PubMed on the genes highly ranked in terms of their Cepo statistics. By querying the gene of interest and the cell type it is assigned to with respect to the Cepo statistics as key terms in the PubMed search, we categorized a gene as either “known” or “new” in terms of the number of publications returned by the query. We considered a gene as “known” if its query returned a search of one or more publications or as “novel” if its query returned no searches (**Figure 3A**). Genes found neither to be a known or novel gene were defined as “others”. We found that whilst many cell identity genes uncovered by Cepo are known gene markers for their respective cell type a large number of genes that were not previously associated with the retinal cell types were ranked highly by their Cepo statistics (**Figure 3B**). Indeed, both known and new cell-type markers uncovered by Cepo showed expression specific to that cell type, whereas the randomly selected stable genes showed non-specific expression across all cell types (**Figure 3C and S3A**). In agreement, known and new gene markers, but not the randomly selected stable genes, showed utility in classifying their respective cell types as indicated by their feature importance score computed from the random forest algorithm (**Figure 3A and S3B**).

**Figure 3.**
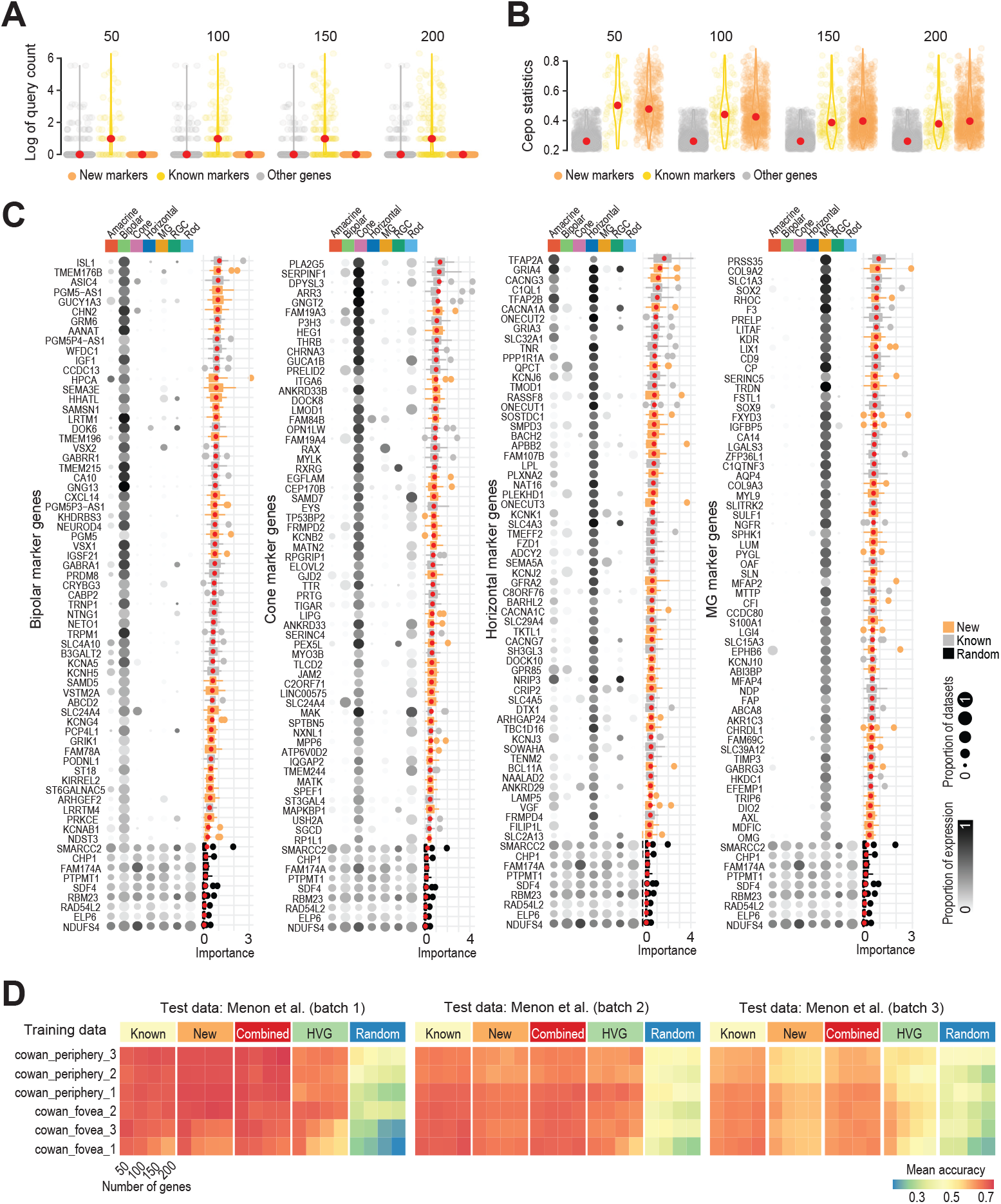
Identification and validation of novel cell-type-specific gene markers of the retina. **(A)** Scatter violin plots of log of query count of the top 50, 100, 150, 200 genes categorized into known or new genes. The scatter plot visualizes the PubMed queries for the results from all cell types and those from non-marker-genes for comparison. **(B)** Scatter violin plots of the same query results as in (A) but of the respective Cepo statistics. **(C)** Cell-type-specific gene markers identified by Cepo. Proportion of cells expressing each marker in each cell type is represented by the gradient color and the proportion of datasets having each marker expressed is represented by the size of the balloons. Importance scores of gene markers are derived from random forest classification of cells using these markers. Novel markers are highlighted in orange and known markers are in gray. Randomly selected genes (in black) are included as control. **(D)** Classification accuracy of an independent test data (Menon et al., 2019) from *k*NN classifiers trained on each of the Cowan datasets using known or new gene markers or their combination, highly variable genes (HVG), and randomly selected genes.

To further evaluate the utility of known and novel cell identity genes in delineating the retinal cell types, we performed cell-type classification by training a *k*-nearest neighbor (*k*NN) classifier using the top genes ranked by the Cepo statistics and classifying an independent retina scRNA-seq dataset generated from Menon et al. (Menon et al., 2019) (**Figure 3D and S3B**). To evaluate the stability of the classification results, we performed the classification analysis on a varying number of top genes (from 50 to 200 genes per cell type). We also assessed the classification accuracy using highly variable and randomly selected genes. We found that the known and new marker genes led to similar classification accuracy of cell types with an average > 0.8. As expected, highly variable genes showed better than random classification accuracy on cell types but were much lower than Cepo selected cell identity genes. Lastly, the combination of known and new marker genes resulted in the best classification in most cases across all three batches of the data from Menon et al. (**Figure S3B**). Taken together, these analyses support the discovery of new gene markers for each of all major retinal cell types and demonstrate their efficacy in delineating their respective cell types from other retina cell types.

### Identifying genes associated with human retina maturation

Several recent studies have profiled the developing human eye using scRNA-seq (Cao et al., 2020; Lu et al., 2020). To characterize these fetal samples and identify genes that are associated with maturation of each retina cell type (**Figure 4A**), we curated these datasets each profiling a wide range of developmental stages (**Figure S4A**) and combined this resource with the mature retinal atlas. After embedding all the single cells into a common space, their visualization on UMAP revealed that the major cell types of the retina clustered together (**Figure 4B and S4B**). In line with the developmental birth order of retinal neurons and the Müller glial, we observed that fetal samples from up to approximately 100 days of conception contained high proportions of retinal ganglion cells, horizontal cells, and cones (in relation to rod cells) (**Figure S4C**) (Shiau et al., 2021). Rods, amacrine, bipolar, and Müller cells demonstrated increased proportions after the 100-day time point, in agreement with the knowledge that these cell types are late-born cells. The final proportions of the fetal and mature cells in the combined retinal atlas demonstrate the inclusion of both age groups in all the cell types (**Figure S4D**).

**Figure 4.**
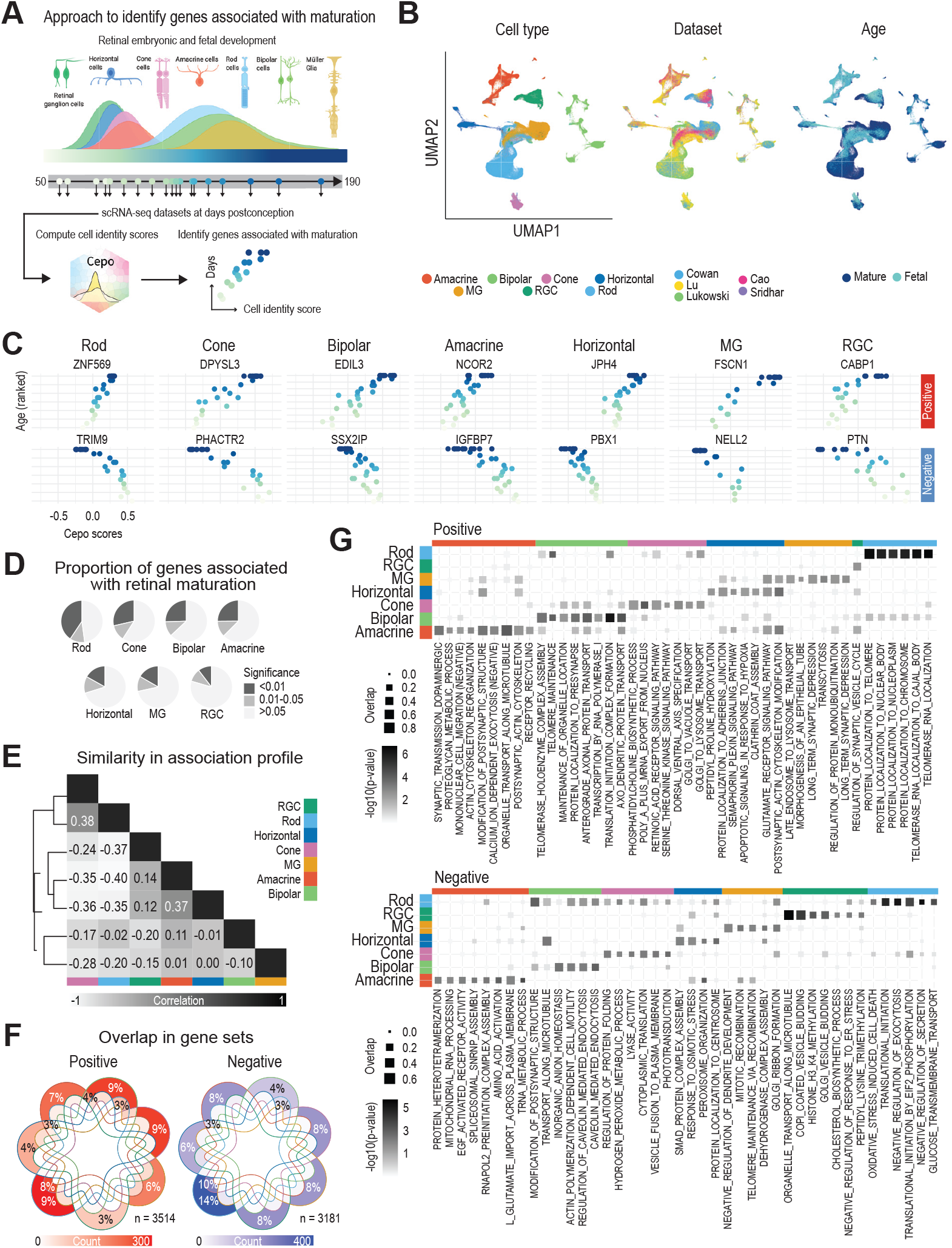
The retinal cell identity scores uncover genes associated with retinal maturation. **(A)** Schematic illustrating the workflow to derive cell-type-specific maturation-associated genes. **(B)** UMAP representation of transcriptome profiles of cells combining both adult and fetal tissues. Cells are coloured by their types (left), datasets (middle), and sample types (right). **(C)** Scatter plot of developmental age (x-axis) and Cepo statistics (y-axis). Points denote individual samples and are colored by their ranked developmental time point. The top panel shows top maturation genes that are positively associated with age and the bottom panel shows those that are negatively associated with age. **(D)** Proportion of highly significant, significant, insignificant maturation-associated genes for each cell type. Highly significant, FDR-adjusted p-value < 0.01; significant, FDR-adjusted p-value between 0.01 and 0.05; and insignificant, FDR-adjusted p-value > 0.05. **(E)** Similarity in maturation association profile between the cell types in terms of the Pearson’s correlation coefficient. **(F)** Venn diagram illustrating the overlap among the positively (left) and negatively (right) significant genes (FDR-adjusted p-value < 0.05). The total percentage of overlap is highlighted for intersections greater than 2% of overlaps. The color scale denotes the absolute number of genes in each gene set. **(G)** Bar plot of -log10(p-value) indicating the significance of enrichment of gene sets.

To discover genes associated with human retinal maturation, we investigated the increase or decrease in their expression specificity with respect to a cell type over time. To this end, for each gene we performed a test of monotonicity of the association between its cell-type-specific Cepo statistics and the developmental age of the retinal samples. This analysis therefore enables the generation of cell-type-specific scores that denote whether a gene exhibits a gain or loss in cell-type-specific expression over time (**Figure 4C**). Cepo statistics were computed as previously described for each cell type and batch and the clustering results from their pairwise correlation relationships visualized as a heatmap (**Figure S4E**). We found that the cell types were associated with varying numbers and proportions of maturation-related genes (**Figure 4D, S4F**). The photoreceptor cells, in particular rod cells, exhibited the highest proportion of genes associated with maturation, with more than half the genes demonstrating a significant association with age. This suggests that photoreceptor cells show a high degree of rewiring during development. The ganglion cells have the least number of genes associated with maturation. As the ganglion cells are the first to be generated amongst the retinal cell types during development (O’Hara-Wright and Gonzalez-Cordero, 2020), this finding suggests that these cells have reached maturation prior to the time course profiled by the fetal scRNA-seq datasets (**Figure S4C**). Furthermore, we found that these maturation profiles were largely cell-type-specific (**Figure 4E**,**F**) and that most of the maturation-associated genes exhibited a gain in cell-type-specificity with age (**Figure S4F**). Collectively, these findings reveal that substantive rewiring occurs during development where most of the rewiring leads to an increase in cell-type-specific gene expression.

Next, we addressed whether the genes associated with maturation of the major cell types of the retina represented those relevant to their respective cell types. To test this, we performed over-representation analyses using the gene sets shown in **Figure S4F**. Consistent with the neuronal identity of many of the retinal cell types, we observed that many of the most enriched pathways among the gene sets positively associated with retinal maturation were related to the formation and regulation of the synapses (**Figure 4G**). For example, the maturation of rods, cones, bipolar, and ganglion was associated with terms such as “regulation synaptic vesicle priming”, “protein localisation to presynapse”, and “regulation of synaptic vesicle cycle”. Horizontal and amacrine cells receive input from many photoreceptors and bipolar/ganglion cells, respectively. In line with their role in receiving and integrating signaling from many populations of cells, one of the most significant pathways enriched during the horizontal and amacrine cell maturation was “modification of postsynaptic structure”. Furthermore, consistent with the fact that amacrine cells are the dopaminergic neurons of the eye, a positive amacrine maturation profile was strongly enriched for the “Regulation of synaptic transmission dopaminergic” pathway (Dacey, 1990). Furthermore, the neurotransmitter that is responsible for the vertical pathways through the retina is glutamate. All photoreceptor cells, bipolar cells, and the retinal ganglion cells therefore release the excitatory amino acid glutamate to transmit signals. Cell types such as the Müller cells, amacrine, and horizontal cells are depolarised by the release of glutamate, through the excitatory synaptic transmission of signal through the glutamate receptors. In line with the importance of this signaling pathway to enable relay of signal, one of the most significant pathways among Müller cells, amacrine, and horizontal cells was “regulation of glutamate receptor signaling pathway”.

### Assessing the fidelity of human retinal organoids generated from diverse differentiation protocols

Recent advances have led to the development of several state-of-the-art protocols for generating human retinal organoids that largely resembles the endogenous retina (**Figure 5 and Table S2**) (Afanasyeva et al., 2021), and several studies have begun profiling these organoids at single-cell resolution and across a range of maturation stages. We curated these public datasets obtained from guided and unguided and 2D-3D and 3D-only protocols, all of which varied in media composition and addition of supplements or growth factors (Cowan et al., 2020; Kallman et al., 2020; Lu et al., 2020; Sridhar et al., 2020) (**Figure 6A and S5**). Whilst functional and molecular studies have evaluated the efficacy of these organoid protocols for efficient and robust generation of retinal cell types, no studies have systematically evaluated these protocols by comparing their global cellular and molecular profiles with human retina tissue. To address this gap, we devised a framework to systematically assess these retinal organoid protocols. In brief, leveraging the mature and fetal retinal atlas, we derived references denoting the cell identity scores, the cell-type proportions, and the cell-type coverage aspired for the organoids. Importantly, we implemented four metrics based on the retinal atlas that measure the capacity of the protocols to mimic the cellular identities of 1) the mature and 2) the fetal retinal tissue, 3) to generate the cell-type proportions found in the tissue, and finally 4) to generate all the major cell types in the retina (**Figure 6B**). The full description of how the references were generated and the metrics are computed can be found in the Methods section.

**Figure 5.**
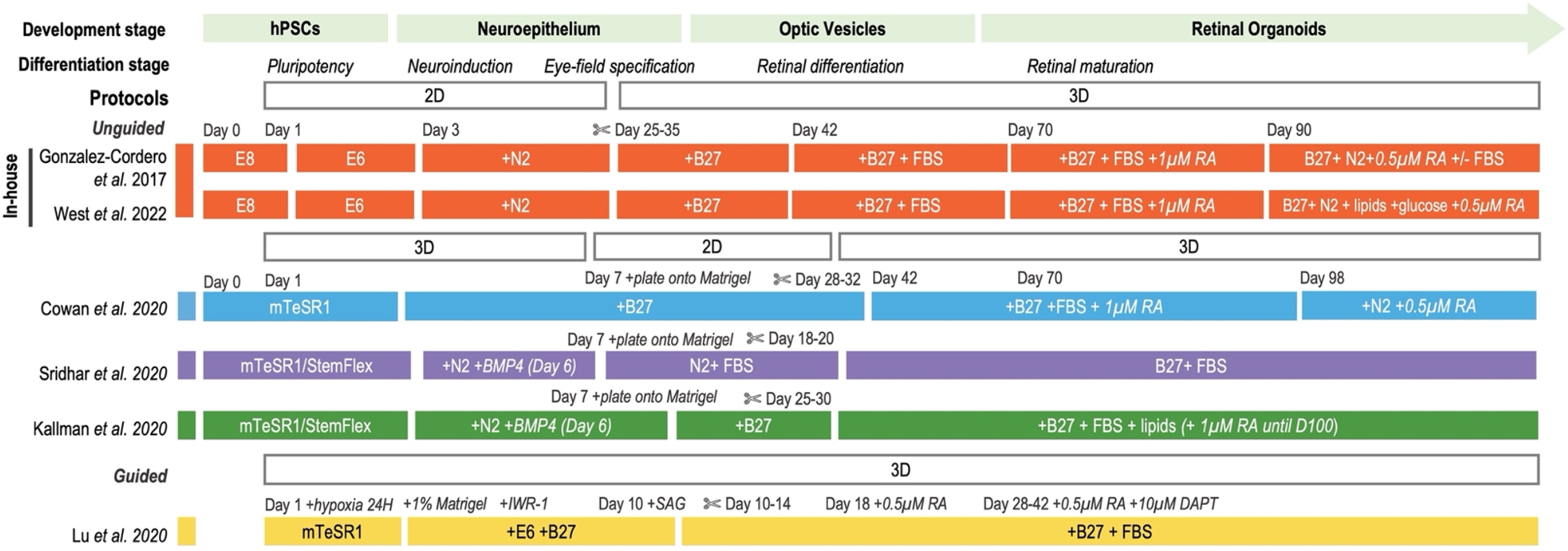
Protocols of human retinal organoids. Schematic view of the culture systems for generating human retinal organoids. The schematic illustrates the timeline of retinal induction, differentiation, and maturation steps, outlining 2D and 3D stages, key supplements and factors added to culture conditions across the published protocols. Abbreviations: hPSCs= human pluripotent stem cells; E8= Essential 8; E6= Essential 6; RA= retinoic acid; H= hour, DAPT= N-[N-(3, 5-difluorophenacetyl)-l-alanyl]-s-phenylglycinet-butyl ester) γ-secretase inhibitor; ?= dissection/dislodgement.

**Figure 6.**
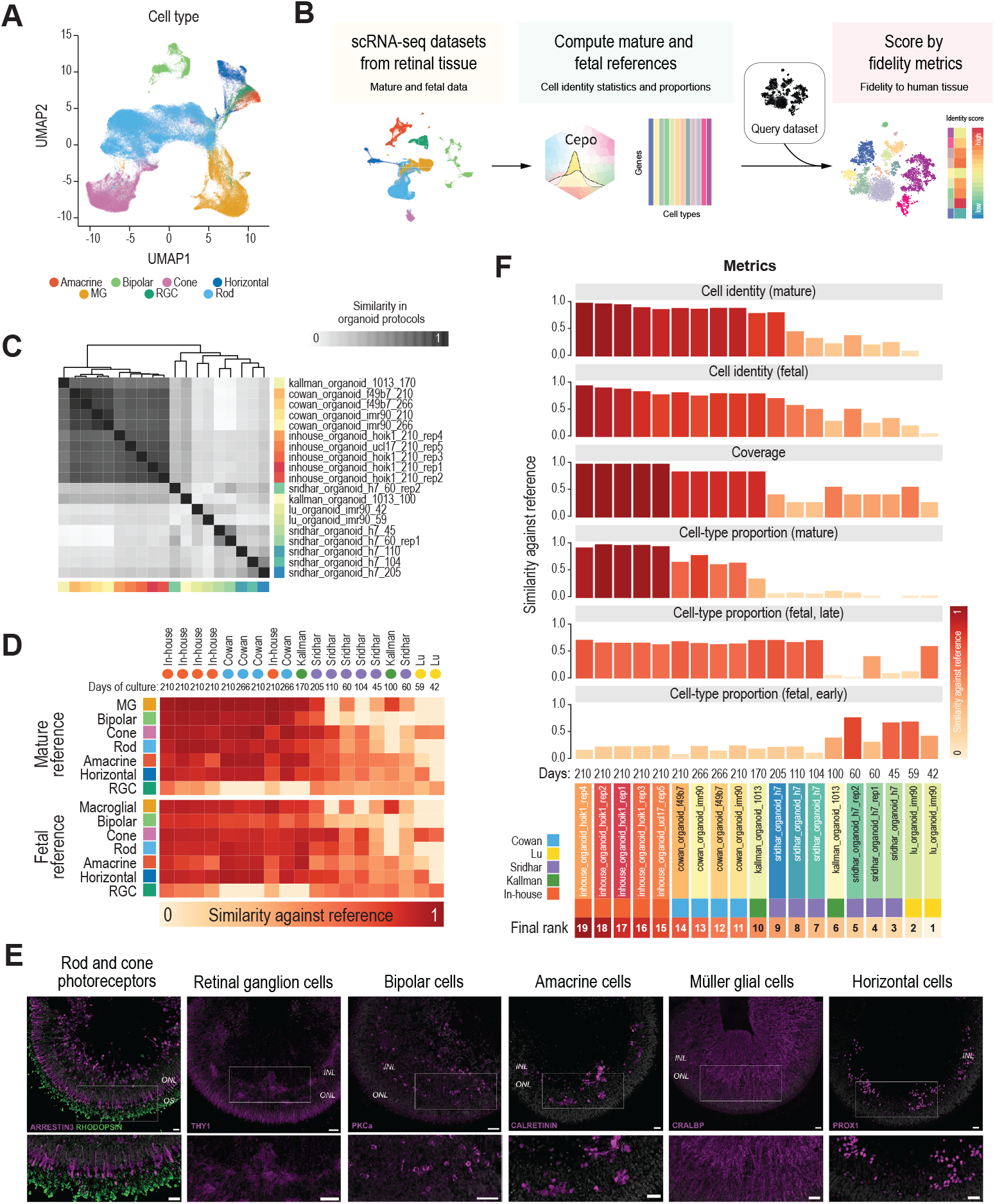
Benchmarking the human retinal organoid protocols in terms of their fidelity to the retina tissue. **(A)** UMAP representation of transcriptome profiles of cells from human retinal organoids. Cells are coloured by their types. **(B)** Schematic overview of the benchmarking procedure and evaluation metrics. **(C)** Similarity in cell identity profiles between the scRNA-seq datasets generated from various organoid protocols. Protocols are color-coded as in (A). Similarity is measured in terms of the Pearson’s correlation coefficient. **(D)** Benchmarking results of the fidelity of the individual cell types to either the mature or fetal reference. The cell identity metric gives a score between 0 and 1, where 1 denotes a high capacity and 1 denotes a low capacity of the protocol to generate cell types that closely match their tissue counterparts. **(E)** Representative images of retinal cell types in human retinal organoids. Abbreviations: ONL:outer nuclear layer, INL: inner nuclear layer, OS: outer segments. Scale bars: 20um. **(F)** Final benchmarking results of the human retinal organoid protocols ranked in terms of the combined score from four evaluation metrics: cell identity (mature), cell identity (fetal), coverage, and cell-type proportion (mature). The scores have been scaled from 0 to 1 within each evaluation metric, where a value of 1 denotes a high capacity of the protocol to mimic the reference and 0 denotes low capacity.

Using this benchmarking framework, we evaluated the capacity of each protocol to generate cell types that closely mimic the human eye. To assess the reproducibility of the protocols, we evaluated the batches within each dataset independently, which led to 2-6 replicates per protocol. We first evaluated the similarity between the protocols by comparing the cell-type identity scores generated for each dataset and batch. Here, we performed scRNA-seq on human retinal organoids cultured for approximately 210 days (30-weeks-old) generating using our own protocol (**Figure 5, S5A**). We found that our in-house generated retinal organoids (Gonzalez-Cordero et al., 2017; West et al., 2022) and those of Cowan et al. demonstrated the highest degree of similarity and also the highest level of consistency between batches (**Figure 6C**). Other protocols (Kallman et al., 2020; Lu et al., 2020; Sridhar et al., 2020) showed more variability between batches and showed distinct cell identity patterns (**Figure 6C**).

Next, for each of the datasets we computed the cell identity metric to assess the fidelity of the individual cell types of the organoids against the mature and fetal references. Ordering the samples by their similarity to the reference revealed that the organoids derived from our in-house protocol, closely followed by the Cowan protocol, consistently outperformed those at similar ages derived from other protocols in terms of achieving a high fidelity in cellular identity against both references (**Figure 6D**). Our immunohistochemistry analysis indeed confirmed the presence of all the major retinal cell types in day 210 human retinal organoids (**Figure 6E**, n=15 organoids; N=3 differentiation batches). Conversely, many of the organoids, particularly those from earlier ages (less than 130 days old) ranked poorly against both the mature and fetal reference and also in terms of coverage, lacking many cell types such as the amacrine and horizontal cells. This may be because retinal organoid maturation follows a three-stage process, wherein each stage progressively leads to a more complex cell composition in the next stage (Afanasyeva et al., 2021). We observed that these organoids were less complex in their cell-type composition and the ganglion cells exhibited the highest similarity to the fetal reference and generally comprised the largest proportion in these earlier time point organoids (**Figure 6E and S5B**). This is consistent with this knowledge and that the retinal ganglion cells are the first to develop in retinogenesis. Furthermore, younger organoids showed greater similarity to the fetal reference than the mature reference. Our benchmarking analyses revealed that human retinal organoids cultured for more than approximately 170 days were more representative of the mature human retina in line with our expectations of organoid maturation.

To perform cross-laboratory and cross-protocol benchmarking of the human retinal organoids, we applied the three fidelity metrics described in **Figure 6F** to the organoid datasets. The cell identity metric was computed as the average of the cell-type-specific scores described in **Figure 6B**. As per their individual scores, the overall score showed that the our protocol generated retinal cell types most fidel to both mature and fetal tissue, closely followed by those from the Cowan protocol, consistent with what were observed in **Figure 6D**. The organoids from the Kallman and Sridhar protocols at similar culture periods (grown until 170 and 205 days, respectively) performed less in terms of the cell identity metrics. Whilst it is not appropriate to directly compare the organoids from the early and late culture stages, it is worth noting that these organoids show minimal resemblance to the fetal reference, suggesting either that progenitor cells present at these early timepoints of organoid are not captured in our fetal reference or that these organoids do not robustly recapitulate the fetal retinal cell types.

Next, the coverage metric was computed as the proportion of the major cell types of the retina present in the organoid data. We observed that our protocol can achieve the highest coverage of the major cell types of the retina consistently across five batches that differ in organoid cell line and organoid batch. (**Figure 6F**). Most of the other protocols (Cowan and Kallman) can generate the majority of the cell types, except retinal ganglion cells, which are known to decrease in numbers with extended organoid cultures. The final metric–the proportion of cell types–was computed from the mature and fetal reference, the latter which is also subdivided into two references denoting early and late fetal maturation. We found that again our protocol excelled the most among the protocols in generating cell-type proportions that represent those found in the mature human eye and not the fetal eye (**Figure 6F**). Intriguingly, even though the early-stage organoids poorly recapitulated the cellular identities of the fetal tissue, we found that their cell-type proportions better reflected those of the fetal eye (**Figure 6F**). Overall, our benchmarking study revealed our in-house protocol generates cell types of the retina with the highest fidelity to the human adult retina in terms of their cellular identities, cell-type proportion and coverage. Lastly, to facilitate future benchmarking of PSC-derived human retinal organoids generated from various differentiation protocols, we have implemented our retinal cell type identity map and the benchmarking framework as an online resource (https://shiny.maths.usyd.edu.au/Eikon/) where the fidelity and quality of organoids can be explored, assessed and scored.

## Discussion

One of the key innovations of our current study is the discovery of potential novel markers of the major retinal cell types and of retinogenesis. In particular, to elucidate this array of cell-type-specific markers, we integratively analyzed the scRNA-seq datasets of the fetal and the mature retina. This is now possible with the advent of single-cell technologies which has enabled the extensive exploration of retinogenesis at a single-cell resolution. For example, recent studies have begun elucidating the trajectories of retinal-derived photoreceptor differentiation, highlighting markers that distinguish rod versus cone specification during development (Kallman et al., 2020). Our study takes advantage of the rich resource of scRNA-seq datasets of the developing and mature retina to interrogate genes that are associated with retinogenesis. We uncovered intrinsic differences in the maturation profile between all the major cell-types of the retina and showed that these specification profiles were largely cell-type specific. Whilst we anticipate that these genes can be used as a resource for further studies to investigate retinogenesis in development and disease, we note that future studies are required to validate these findings in the *in vivo* and *in vitro* systems.

Another innovation is our comprehensive effort to benchmark human retinal organoids generated by diverse differentiation protocols against the *in vivo* human retina. Our benchmarking study revealed that unguided human retinal organoids generated using our in-house differentiation protocol and that of the Cowan et al. were top performers (Cowan et al., 2020; Gonzalez-Cordero et al., 2017; West et al., 2022). Notably, these two protocols have similar time-course of RA addition and supplementation. While these methods considered the maturation stages of development and made suitable modifications at late stages in culture, Sridhar and Kallman protocols maintained organoids in more basic retinal differentiation media throughout. All differentiation protocols showed efficient generation of organoids, however our 2D-3D is able to generate organoids from PSC confluence in a simpler workflow (**Figure 5**) (Gonzalez-Cordero et al., 2017; Reichman et al., 2014; West et al., 2022).

Furthermore, we have recently shown that optimisation of late-stage culture conditions with lipids supplementation enhances maturation of photoreceptor cells (West et al., 2022). Amongst the benchmarked protocols, our protocol generates cell types of the retina with the highest fidelity to the human retina tissue across several metrics: their cellular identities, cell-type proportions and coverage (**Figure 6D-F**). Our benchmarking highlights that the choice of retinal differentiation protocols is an important consideration when using human retinal organoids as models of disease and as tools to establish efficacy for new therapies.

Whilst our benchmarking study highlighted that some of the protocols are able to successfully develop retinal organoids closely resembling many aspects of the human retina, it also highlighted remaining challenges. We observe that the organoids, in particular those collected at earlier maturation stages (50-170 days), do not fully recapitulate the developmental stages of the human retina (**Figures 6D, F**): a feature to be taken into consideration when investigating human eye development and retinogenesis using human retinal organoids. Independent of the type of differentiation protocols, the development of retinal cells in human retinal organoids follow three stages of development: in the first stage (approximately 30 days of culture), the neuroblastic and retinal ganglion cell layers have formed; in the second stage (approximately 120 days of culture), the outer and inner nuclear cell layers appear; and in the final and third stage (approximately 180 days of culture), the photoreceptor cells and the outer plexiform layers have formed. By 200 days of culture, these retinal organoids are thought to have fully developed into mature cell types representing those of the retina. Further scoring of early-stage organoids using scRNA-seq datasets as inputs into Eikon will offer further insight into the gold standard protocols at early time points of development; therefore, we encourage researchers with such data to use Eikon to assess the fidelity of their retinal organoids.

One of the key limitations of the study is that the fidelity metrics do not address important features of the retina such as the profiles of the other omic layers, spatial patterning, and retinal organoid functionality. Recently, a few studies have begun profiling the retinal organoid using single-cell technologies other than the transcriptomes (e.g., scATAC-seq). The study by Thomas et al. have mapped the cis-regulatory elements of the developing and mature human retina and showed that the human retinal organoids are capable of emulating the DNA accessibility of the human retina (Thomas et al., 2022). The spatial organization of the retina is an important component to evaluate for tissues like the retina that have highly complex, ordered, and specialized structures. Towards this end, spatial transcriptomics that can profile the transcriptomes whilst retaining the spatial coordinates of the single cells would enable us to assess the capacity of protocols to generate organoids that are spatially organized to closely emulate that of the *in vivo* tissue (Rao et al., 2021). Future studies will be required to develop computational methods that can quantify the reference spatial organization for each cell type of the retina and measure the fidelity of organoids to conform to the reference. Lastly, our study did not look at retinal organoid functionalities as one of the fidelity metrics. Several important functions of the eye, including the visual cycle, retinal ON/OFF pathways, and synaptogenesis, were not incorporated (Fathi et al., 2021). Yet these functional studies are paramount to evaluate that the retinal organoids not only look like the retina but also function as such.

Technologies like single-cell patch clamp recordings can identify the hyperpolarization and depolarization of photoreceptors in the organoids, providing a measure of how well cells respond to light and darkness, respectively (Zhong et al., 2014). A recent study from Saha et al. (Saha et al., 2022), demonstrated wavelength-specific light-evoked responses from organoid-derived cone photoreceptor cells similar to electric responses observed in the primate fovea. Finally, cell replacement studies demonstrated the restoration of visual function in advanced disease after transplantation of a purified population of PSC-derived cone photoreceptors isolated from our human retinal organoids (Gonzalez-Cordero et al., 2017; Ribeiro et al., 2021). Therefore, incorporating other omics, spatial, and functional information, along with the transcriptome, will in the future enable us to comprehensively evaluate retinal organoids in health, development, and disease.

## EXPERIMENTAL PROCEDURES

### EXPERIMENTAL MODEL AND SUBJECT DETAILS

#### Human induced pluripotent stem cells

HPSI0314i-hoik_1 (RRID:CVCL_AE82) was obtained from ECCAC. UCLOOi017-A-1 was derived from healthy donor peripheral blood mononuclear cells (PBMCs) as described previously (Fernando *et al*.).

## METHOD DETAILS

### Cell culture and retinal organoid generation

#### Human induced pluripotent stem cell maintenance and differentiation

Cells were incubated at 37°C 5% CO^2^. hiPSCs were grown and expanded in feeder-free conditions using Essential 8 media (E8, Life Technologies) on 6 well plates coated with Geltrex (Invitrogen) at a concentration of 1:100. Media was replaced daily and cells were passaged at 70% confluency via 5-10 minute 37°C incubation with Versene Solution (0.48 mM) (Life Technologies) to detach clumps of cells. Cell clumps were resuspended at a ratio of 1:6-1:12 in E8 with 10µM ROCK inhibitor (Y-27632 dihydrochloride, Tocris) and seeded in fresh Geltrex-coated 6 well plates. For differentiation, hiPSCs were grown to 90-100% confluency.

#### Generation of retinal organoids from hiPSCs

Retinal organoids were differentiated as previously described (Gonzalez-Cordero et al., 2017; West et al., 2022) with some modifications. Briefly, at 90-100% confluency (denotated day 1), hiPSC media was replaced with Essential 6 (E6, Life Technologies) daily for two consecutive days. On day 3, E6 media was replaced with pro-neural induction media (PIM, Advanced DMEM/F12, 1X N2 supplement, 1.9 mM L-glutamine, 1X MEM-NEAA, 10% Antibiotic-Antimycotic, [all Life Technologies]). Optic vesicles displaying neuro-retinal epithelium were manually isolated using a needle under EVOS XL microscope (Invitrogen) between day 25-35 and transitioned to 3D suspension culture in low-binding 96 well U-shaped plates and retinal differentiation media (RDM, DMEM high glucose 68% v/v, Ham’s F-12 Nutrient Mix with GlutaMAX Supplement 29% v/v, 1X B-27 Supplement minus Vitamin A, 10% Antibiotic-Antimycotic [All Life Technologies]). At day 42, RDM was replaced with RDM + Factors (RDMF, DMEM high glucose 60% v/v, Ham’s F-12 Nutrient Mix with GlutaMAX Supplement 26% v/v, 2X GlutaMAX Supplement, 1X B-27 Supplement minus vitamin A, 10% Antibiotic-Antimycotic) [All Life Technologies], FBS 10% v/v [Bovogen]. At day 70, retinal organoids were transferred into low-binding 24 well plates and media was replaced with ALT70 (Advanced DMEM/F-12 85% v/v, 10% FBS, 2X GlutaMAX Supplement, 1X B-27 Supplement minus Vitamin A, 10% Antibiotic-Antimycotic, [All Life Technologies} 100 µM taurine [Sigma Aldrich]) and supplemented with 1uM all-trans retinoic acid (RA) to enhance photoreceptor development. At day 90 and until experimental end-point, media was replaced with ALT90 (Advanced DMEM/F-12, 2X GlutaMAX Supplement, 1X B-27 Supplement minus Vitamin A, 1X N2 Supplement, 7 mM glucose, 10% Antibiotic-Antimycotic, 1X Lipid Mixture, [All Life Technologies] 100 µM taurine [Sigma Aldrich], +/- 10% FBS [Bovogen]) and supplemented with 0.5 uM RA. Media was replaced Monday, Wednesday, Friday and cells maintained at 37°C 5% CO^2^.

#### Immunohistochemistry

Retinal organoids were washed with PBS, fixed for 40-60 minutes in 4% paraformaldehyde prior to incubation in 20% sucrose. Organoids were embedded in OCT, frozen in liquid nitrogen, and then cryo-sectioned at 14 µm thickness. Cryosections were blocked in 5% serum in blocking solution (1% bovine serum albumin in PBS with 0.1% Triton-X) for 2 hours. Primary antibody (**Table S3**) diluted in the blocking solution was incubated overnight at 4°C. Sections were washed with PBS and incubated with secondary antibody (Alexa fluor 488, 546 secondary antibodies) at room temperature for 2 hours. Sections were counterstained with DAPI.

### Single-cell RNA sequencing of retinal organoids and human retina

#### Dissociation of organoids into single cells

Five independent organoid batches were derived for both HPSI0314i-hoik_1 and UCLOOi017-A-1 hiPSC lines. One organoid was dissociated per sample. Retinal organoids were dissociated into single cell suspension using the Neurosphere Dissociation Kit (P) (Miltenyi Biotec). Enzymatic digestion was performed as per manufacturer protocol for 10 minutes at 37°C with intermittent agitation, followed by gentle mechanical dissociation with a p1000 pipette, and a further 5 minute 37°C incubation. The cell suspension was passed through a p200 to ensure single cell dissociation before the enzymatic reaction was stopped by washing with HBSS. The cell suspension was filtered through MACS SmartStrainer 30µm (Miltenyi Biotec) before being pelleted by centrifugation at 400*g* for 10 minutes at room temperature. The cell pellet was resuspended in ALT90 and maintained on ice.

#### Single cell RNA-sequencing

Each single cell suspension of dissociated retinal organoid was assessed for viability using 0.4% Trypan Blue staining on a Countess II Automated Cell Counter (Invitrogen) and concentration was adjusted to 1000 cells/µl. Cell suspension was loaded on a single-cell-B Chip (10X Genomics) for a target output of 10,000 cells per sample. Single-cell droplet capture was performed on the Chromium Controller (10X Genomics). cDNA library preparation was performed in accordance with the Single-Cell 3’ v3 protocol. Libraries were evaluated for fragment size and concentration using Agilent HSD5000 ScreenTape System. Samples were sequenced on an Illumina NovaSeq6000 instrument according to manufacturer’s instructions (Illumina). Sequencing was carried out using 2×150 paired-end (PE) configuration with a sequencing depth of 40,000 reads per cell. The sequences were processed by GENEWIZ, China.

## QUANTIFICATION AND STATISTICAL ANALYSIS

### Public single-cell RNA-seq datasets

#### Data collection

The raw gene-cell count matrices of scRNA-seq datasets were retrieved from the NCBI Gene Expression Omnibus (GEO) unless otherwise stated. A summary of the public datasets used in this study is provided in **Table S1**. The datasets are all related to the human retina, originating either from tissue or organoid and from mature or immature cells.

#### Processing of scRNA-seq and filtering of cell types and suboptimal samples

All scRNA-seq libraries from human retinal tissue and organoids were integrated by Seurat (v4.0) (Hao et al., 2021) under the R (v4.1.1) environment. For each of the published datasets, the cell type labels from the original study were used. Unless specifically specified as suboptimal cells in the metadata, we considered all cells assigned to a cell type as cells that passed the quality control in the original study.

#### Derivation of cell identity gene statistics and assessment of their concordance across batch *Computation of cell identity gene statistics*

To derive the cell identity gene statistics for the human retina atlas, the count matrix of cell-gene variables were first log-transformed and normalized using the logNormCounts function from the scater package (McCarthy et al., 2017). The transformed and normalized data from each batch were subsequently analyzed using the Cepo package (Kim et al., 2021) for quantifying cell identity gene statistics for each major cell type based on the differential stability metric. For comparison, alternative methods (e.g., limma (Ritchie et al., 2015), voom (Law et al., 2014), edgeR (Robinson et al., 2010)) based on differential expression analysis were also used for calculating cell-type-specific gene statistics.

#### Clustering of cell identity gene statistics and evaluation of batch mixing

To evaluate the consistency of the cell identity gene statistics derived for each cell type, clustering was performed on these statistics derived from different batches and datasets and then the degree of batch mixing was determined for three sources of variation: cell type, dataset source, and batch source. Ideally, cell identity gene statistics should reflect the biological identity of the cells and therefore the source of variation should solely arise from cell type identity and not from batch or dataset source.

To generate the cluster, three clustering algorithms (hierarchical clustering, *k*-means clustering, and Louvain clustering were performed. For each of the sources of variation, the resolution of the hierarchical and *k*-means clustering was set by controlling the parameter, number of trees or *centers*, respectively, to equal the number of cell types (7), the number of datasets (3), or the number of batches (14) present in the comparison. For the Louvain clustering wherein setting the exact cluster number is infeasible, the resolution was controlled by setting differing values of *k* when building the shared nearest neighbor graph (cell type, *k* = 5; dataset, *k* = 8; and batch, *k* = 2).

To assess the clustering performance of Cepo-derived cell identity gene statistics, we used the adjusted rand index (ARI) or the normalized mutual information (NMI) to evaluate the concordance of clustering results with respect to the cell type labels, batch, and dataset source, denoted respectively as either *ARI*_*cellTypes*_, *ARI*_*dataset*_, and *ARI*_*batch*_ or *NMI*_*cellTypes*_, *NMI*_*dataset*_, and *NMI*_*batch*_. Considering that the Cepo-derived gene statistics are partitioned into different classes with respect to cell type labels, batch, or dataset source, let - be the number of pairs of samples partitioned into the same class by a clustering method, *b* be the number of pairs of samples partitioned into the same cluster but in fact belong to different classes, *c* be the number of pairs of samples partitioned into different clusters but belong to the same class, and *d* be the number of pairs of samples from different classes partitioned into different clusters. Then the ARI is calculated as follows:

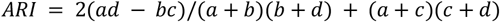

Considering that the Cepo statistics are partitioned into different classes with respect to cell type labels, batch, or dataset source, let *Y* be the clustering outcome by a clustering method and *C* be the original labels of the different classes. Given that *Y* and *C* are the partitions of the same data, the overlaps between the two random variables can be counted and represented as a contingency table. Using information theory to measure agreement between the partitions and maximum likelihood estimation, the empirical joint distributions of clusterings *Y* and *C* are measured. Therefore, using the probabilities that an element falls into a given cluster, the entropy for clusterings *Y* and *C, H*(*Y*) and

*H*(*C*), respectively, and the mutual information *I*(*Y*; *C*) can be calculated. Then the NMI is calculated as follows:

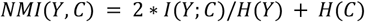

#### Evaluation of concordance of cell identity gene statistics between batches

The evaluation of scRNA-seq datasets from multiple sources oftentimes requires technical noise to be removed. This so-called “batch effect” has been widely addressed through the development of several algorithms (Tran et al., 2020). However, the danger of overcorrection and restriction to a subset of features during the correction has hampered the widespread use of batch correction methods for downstream analysis. Therefore, an approach that bypasses the need to remove batch effects, does not compromise the biological signal, and retains many of the features is ideal to enable a comprehensive comparison between multiple atlases, often with a high degree of sparsity.

To this end, we assessed the fidelity of cell identity gene statistics between varying degrees of batch effect to determine the comparability of Cepo statistics across independent batches. A combinatorial assessment of cellular identity gene statistics concordance was performed between pairs of datasets. For all combinations of samples in the adult retinal tissue atlas, the extent of variance in principal component space contributed by batch and was quantified using the pvca package in R (Bushel, 2021). Briefly, principal variance component analysis fits a mixed linear model using the factor of interest, batch source, as an independent random effect to the selected principal components. Using the eigenvalues as weights, the associated variations of factors are standardized and the magnitude of the source of variability is presented as a proportion of the total variance. Therefore, a greater proportion of total variance denotes the presence of greater batch effect between the dataset pair.

To evaluate the concordance in cell identity gene statistics between the batches, the Cepo statistics were computed for each gene and for each cell type independently within each batch. Then for each batch pair, the concordance of cell identity was measured as the Pearson’s coefficient of the correlation between the Cepo statistics. Finally, the relationship between the concordance and the degree of batch effect was examined. For comparison, cell-type-specific gene statistics from Limma, Voom, and edgeR were also analyzed using the same procedure.

### Identification of novel markers of cell type

#### Categorization of known and unknown cell-type markers

To systematically categorize a predicted marker gene as either already described in the literature or a new cell-type marker, an advanced PubMed query was performed using the R package easyPudMed (Fantini, 2019) for each of the top 500 marker genes. For each cell type, the advanced query consisted of three search terms: 1) the name of the candidate gene of interest; 2) the cell type of interest; 3) and the term “RETINA”, all combined with the operator “AND”. The search terms covered “All Fields” of the possible search items. Any genes for which the PubMed search did not return a publication were categorized as a novel marker gene for the cell type of interest. In contrast, any genes for which the PubMed search did return at least one publication were considered known marker genes for the cell type of interest. One of the limitations of the PubMed query is that the individual publications returned as associated with a gene and cell type combination were not individually assessed for false positive result. Nevertheless, this approach provides a systematic and fast means to screen for novel markers, overcoming the need to manually survey the literature for all combinations of genes and cell types.

#### Quantification of feature importance

To demonstrate the importance of the potential marker genes, we performed feature selection analysis based on the random forest classifier (Breiman, 2001). The Gini index, which measures how important a selected feature is when training the random forest classifier, was used as a proxy for feature importance. To build the random forest classifier, the single-cell transcriptomes were subjected to random stratified sampling to 30% of its original size, and then pseudo-bulk transcriptomes for each cell type were generated by taking the mean expression of the genes. Approximately 350 pseudo-bulk transcriptomes were generated by repeating the sampling procedure 50 times. The random forest classifier was trained using these transcriptomes. As control, 10 cell-type invariant genes, as determined by their Cepo statistics, were included. A separate random forest classifier was built for each batch. Finally, the values for the mean decrease in Gini were extracted from the classifiers and visualized as a boxplot.

#### Classification of test data using known and novel gene markers

To further support the utility of the identified genes as novel markers of their respective cell types, we performed classification of single-cell transcriptomes of the human retina from an independent study (Menon et al., 2019), which was not included in the datasets used to derive the cell identity gene statistics. The *k*-nearest neighbor (*k*NN) classifiers were trained by varying the following four conditions:

##### Training data

Single-cell transcriptomes from six batches of data from the Cowan et al. (2021) study were used. The three batches of single-cell transcriptomes were each derived from either the periphery or the fovea.

##### Testing data

Single-cell transcriptomes from three batches of data from the Menon et al. (2019) study were used. The scRNA-seq data were derived from human retina tissue from either the macula in the central retina or a region of the mid-peripheral retina. Of note, the retina was mechanically separated from the retinal pigment epithelium-choroid.

##### Number of gene markers

The number of gene markers included in the training.

##### Category of gene markers

The top gene markers were derived from five categories: (1) gene markers categorized as known; (2) gene markers categorized as novel; (3) a gene set with both known and new markers; (4) highly variable genes; and (5) randomly selected genes. For gene sets 1-3, the top genes were ordered by Cepo statistics. For gene set 4, the top genes were ordered by the FDR-adjusted p-values computed from fitting a trend on the variance and mean of the log gene expression values.

Finally, the number of neighbors was set to *k* = 3 for all the *k*NN classifiers, and the final classification accuracy is calculated as the accuracy across each cell type *c* as follows:

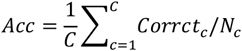

where *C* is the total number of cell types in a dataset, *Corrct*_*c*_ is the correct prediction for cell type *c*, and *N*_*c*_ is the number of cells in cell type *c*.

### Identification of maturation-associated genes in retinal development

#### Determination of genes associated with maturation

To determine the genes associated with maturation, the Cepo statistics derived for all the batches originating from the fetal and mature retinal atlas were used. For each gene, we computed the Spearman correlation coefficient as a function of the change in its Cepo statistics over developmental age. We calculated the monotonicity of the association between the two variables *X* the ranked Cepo statistics and *Y* the ranked developmental age using the following:

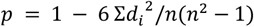

where *p* is the Spearman’s rank correlation coefficient, *d*_*i*_ is the difference between *X* and *Y*, and *n* is the number of observations.

#### Measurement of similarity of maturation-association profiles between cell types

To investigate the similarity in the profile of maturation between cell types, we performed hierarchical clustering on the pairwise correlation matrix of the Spearman coefficient statistics on all the genes found to be significantly associated with maturation (FDR-adjusted p-value < 0.05) in at least one cell type. The similarity is visualized as a clustered heatmap where 1 denotes complete positive correlation and -1 denotes complete negative correlation.

#### Gene set over-representation analysis of maturation-associated genes

Gene set over-representation analysis was performed on the genes significantly associated with maturation. Significance in association was determined as the FDR-adjusted p-value lower than 0.05. The gene set enrichment analysis was performed for each cell type and on either the gene sets positively or negatively associated with maturation. Using the GO terms related to biological processes from the C5 ontology gene set from the MSigDB collection (Liberzon et al., 2011), we assessed the over-representation of these gene sets among the maturation-associated genes using the fgsea R package (Korotkevich et al., 2022).

### Analysis of in-house single-cell RNA sequencing data generated from retinal organoids

#### Read alignment and expression count table generation

From the sequencing results of the 10x Chromium experiments, the unique molecular identifiers, cell barcodes, and the genomic reads were extracted using Cell Ranger with default parameters (v3.1, 10x Genomics). The extracted reads were aligned against the annotated human genome, including the protein and non-coding transcripts (GRCh38, GENCODE v27). The reads with the same cell barcode and unique molecular identifier were collapsed to a unique transcript, generating the count matrix where columns correspond to single cells and rows correspond to transcripts. To remove potentially empty droplets with ambient RNA, the emptyDrops function from the DropletUtils package was used. Droplets with significantly non-ambient profiles were called at a false discovery rate of 1%, applying the Benjamini-Hochberg correction to the Monte Carlo p-values to correct for multiple testing. Next, to remove potentially unhealthy or suboptimal cells, cell filtering was performed using the number of reads, the proportion of genes expressed, and the fraction of mitochondrial reads as criteria. Specifically, as cells with greater than 10,000 reads, 99% of genes not expressed, and 25% of mitochondrial gene expression were removed. Transcripts from mitochondrial- and ribosomal-protein coding genes were discarded for downstream analyses such as embedding and clustering, because they are typically known to be highly expressed irrespective of biological identity.

#### Doublet detection and filtering

The presence of multiplets in single-cell data can arise from incomplete dissociation of single cells meaning that more than one cell can be encapsulated in GEMs. DoubletFinder, an algorithm to detect multiplets in single-cell data, was used to remove potential doublets or multiplets from each biological batch at a threshold of 5.0% (McGinnis et al., 2019).

### Integration and clustering

#### Embedding transcriptomes into a shared latent space

To embed the single-cell transcriptomes into a shared latent space, for each batch the count matrix was first normalized to the total number of reads and then factored by a 10,000 scaling factor. Then the top 2,000 features, among the top 1,000 highly variable features determined through variance stabilizing transformation, were prioritized by their variance across all the scRNA-seq batches. Next, the cell pairwise anchor correspondences between different single-cell transcriptome batches were identified with 30-dimensional spaces from reciprocal principal component analysis (Hao et al., 2021). Using these anchors, the scRNA-seq datasets were integrated and transformed into a shared space. Gene expression values were scaled for each gene across all integrated cells and used for principal component analysis (PCA). For the integration of the organoid datasets, *k. filter* and *k. weight* were set to 160 and 90, respectively, to accommodate the integration of datasets with fewer than 200 cells.

To generate the embeddings containing the single-cell transcriptomes derived from the mature tissue, the single cells were embedded into two-dimensional UMAP space by using the first 15 principal components (PCs). To generate the embeddings containing the single-cell transcriptomes derived from the mature and development tissue combined, the single cells were embedded into two-dimensional UMAP space by using the first 30 principal components (PCs). Finally, to generate the embeddings containing the single-cell transcriptomes derived from the retinal organoids, the single cells were embedded into two-dimensional UMAP space by using the first 15 principal components (PCs).

#### Clustering and classification of in-house datasets

To cluster the single cells from the organoid datasets, the shared nearest neighbor graph was constructed on the first 30 PCs of the shared embedding using the default arguments of the FindNeighbors function in the Seurat package. Then the Louvain clustering algorithm with resolution equal to 1.1 was used to cluster the single cells. Classification of the single cells from the in-house datasets was performed by assigning them to the cell type that according to the labels of the public datasets that most dominate the cluster assigned to the cell of interest.

### Analysis and benchmarking of retinal organoid protocols

#### Similarity of organoid protocols to one another

To evaluate the closeness of the organoid protocols to one another, we first performed Cepo analysis on the individual batches of the organoid datasets as in subsection “Computation of cell identity gene statistics” to derive cell identity gene statistics for each of the major cell types in each batch. Then, intersecting on the genes that are commonly found in all 19 batches, we aggregated the Cepo-derived gene statistics into a single vector. The similarity of cell identity profiles between the batches was evaluated in terms of the Pearson’s correlation coefficient between the aggregated statistics. A hierarchical clustering of the pairwise similarity matrix was performed to cluster like protocols together and discern disparate protocols.

#### Evaluation of the fidelity of organoids to the human tissue

To evaluate the fidelity of organoids to the human tissue, we generated the following five evaluation metrics:

1. **The cell-type-specific similarity against the cell identity retina reference** The cell-type-specific similarity of the query organoid against the cell identity retina reference resolved by mature and developmental cells were computed. The reference consists of the top Cepo statistics for all the major cell types of the mature and development atlas. The average cell identity scores from the mature cell types and the developmental cell types were taken as the reference. Then, to compute the cell-type-specific similarity of the query data, we calculated the Pearson’s correlation coefficient between the Cepo statistics derived for the query data and each of the references for each cell type.
2. **The overall similarity against the cell identity retina reference** The averaged score of the cell-type-specific similarities from (1) were computed for each protocol across cell types to generate the overall cell identity score. As in (1), a higher score denotes stronger fidelity to the retinal reference and a lower score denotes weaker fidelity.
3. **The coverage of cell types** The coverage of cell types generated from the organoid protocol was calculated. The expected retinal cell types are cones, rods, Müller glial, ganglion, bipolar, horizontal, and amacrine cells. Protocols with the capacity to generate all cell types were assigned a score of 1, whereas those with nil capacity to generate the expected cell types were assigned a score of 0.
4. **The concordance in cell-type proportion with the retina reference** The concordance against the proportion of cell types expected in the retina was measured using the averaged proportion profiles of the mature and fetal retina tissue datasets. To account for the differences in cell-type proportions that result from early and late-born cell types in early and late retinogenesis, the fetal proportional reference was sub-categorized into two time points (early [<97 days] and late [>97 days]). The intraclass correlation coefficient (ICC) for oneway models was used to compute an index of consistency of the proportions (Gamer et al.). A protocol with a high ICC reflects a high capacity to reciprocate the proportions found in the real tissue, whereas those with a low ICC reflect those with a low capacity.

For each metric, we then aggregated the results for all the benchmarked protocols and rescaled the score to a range of [0, 1]. Finally, equally weighting these metrics, the average of the overall cell identity (mature and fetal), the coverage, the cell-type proportion (mature), and 1 minus the cell-type proportion (fetal, early) was taken to generate a final score. This score was used to benchmark the retinal organoid protocols in terms of their fidelity to the human retinal tissue.

### Development of the Eikon software

We implemented Eikon, an interactive web tool to facilitate the assessment of the fidelity of retinal organoids to the *in vivo* retinal tissue. Eikon accepts a SingleCellExperiment object (Lun et al., 2022). Multiple parameters can be specified in Eikon, including the assay, age of samples, and whether normalization is required. Users can customize the visualization plots using the provided options and all key visualizations are downloadable. Specifically, Eikon outputs three key visualizations, including several correlation heatmaps, reduced dimension plots, and an interactive table of Cepo statistics for each retinal cell type contained in the query data. The correlation heatmaps display the correlation cell identity scores between the query and reference datasets, allowing users to assess the fidelity of their data in a visually intuitive manner. Importantly, the query data can be compared to all or specific developmental stages of the reference dataset. PCA, UMAP, and t-SNE plots are also displayed and can be coloured by variables contained in the query data such as the proportion of zeroes and UMIs. Additionally, plots can be coloured according to the expression levels of a particular gene of interest which can be found using an interactive table displaying the Cepo statistics for each gene of each retinal cell type in the query data.

## Data and code availability

The accession numbers of public datasets used in this study are listed in **Table S1**. The sequencing data generated in this study has been deposited at Gene Expression Omnibus (GEO) under the accession number GSE201356. The code generated during this study is available upon reasonable request to the Lead Contact. Eikon (https://shiny.maths.usyd.edu.au/Eikon/) is available as an interactive web application to explore the fidelity of retinal organoids.

## Acknowledgments

We thank the intellectual feedback from the Embryology Unit at Children’s Medical Research Institute (CMRI) and engagement from the colleagues at the School of Mathematics and Statistics, The University of Sydney and Sydney Precision Bioinformatics Alliance. We acknowledge the CMRI Single Cell Analytics, Advanced Microscopy Centre and the Australian Cancer Research Foundation Telomere Analysis Centre for supporting this work. This work is funded by National Health and Medical Research Council (NHMRC) Investigator Grant (1173469) to P.Y., Ophthalmic Research Institute of Australia, and Luminesce Alliance (PPM1 K5116/RD274). Luminesce Alliance is a non-profit joint venture between CMRI, the Sydney Children’s Hospitals Network, and the Children’s Cancer Institute and is affiliated with the University of Sydney and the University of NSW. The graphical abstract was generated using BioRender (https://biorender.com).

## Author contributions

Conceptualization, H.J.K., A.G.C., P.Y.; Methodology, H.J.K., M.O-W., A.G.C., P.Y.; Investigation, H.J.K., D.K., A.G.C., P.Y.; Data generation, M.O-W., T.H.L., B.Y.L., R.V.J., A.G.C., P.Y.; Writing – Original Draft, H.J.K., M.O-W., A.G.C., P.Y.; Writing – Review & Editing, all authors; Supervision, R.V.J., A.G.C., P.Y.; Funding Acquisition, R.V.J., A.G.C., P.Y.

