## Supplementary figures for "Comprehensive Characterisation of Fetal and Mature Retinal Cell Identity to Assess the Fidelity of Retinal Organoids"

### Supplementary figures and tables

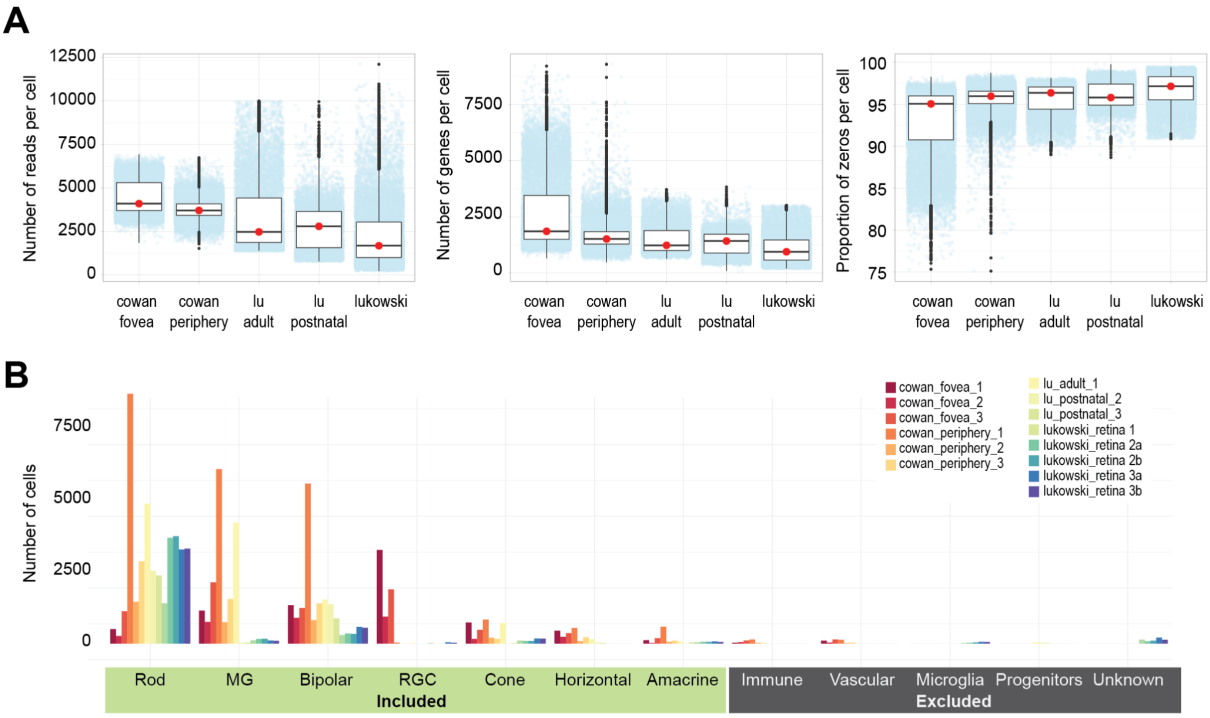


**Figure S1. Quality control of retinal mature tissue scRNA-seq datasets. (A)** Number of reads assigned, genes quantified, and proportion of zeros in each cell for each human retina dataset. **(B)** Number of cells identified for each cell type in each dataset and batch.

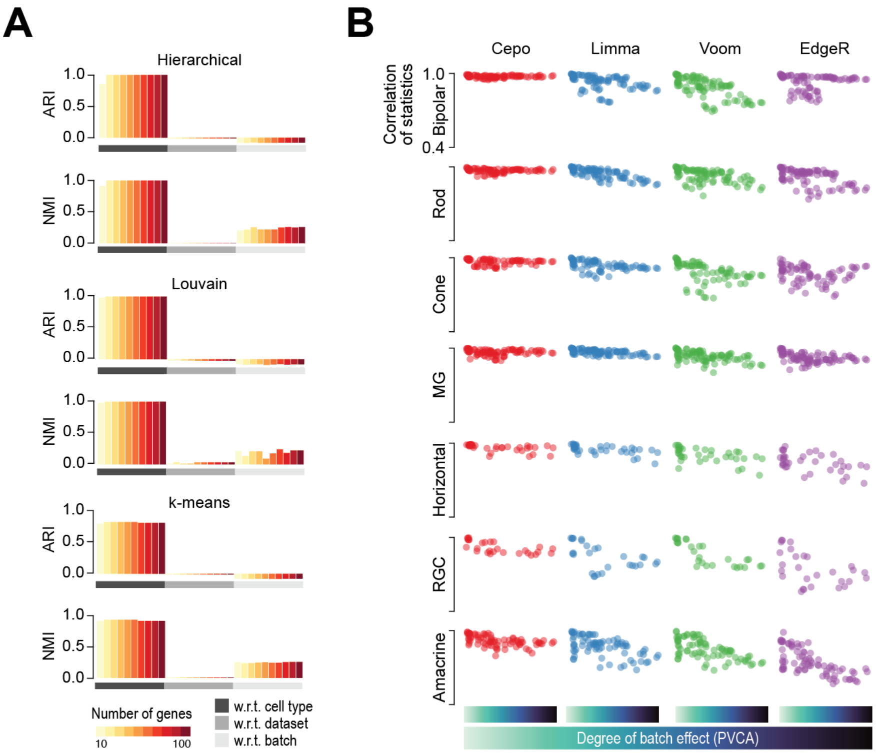


**Figure S2. Assessment of batch effect on deriving cell-type-specific gene statistics. (A)** Clustering concordance quantified by adjusted Rand index (ARI) and normalized mutual information (NMI) with respect to cell type, dataset, and batch using hierarchical, Louvain, and k-means clustering algorithms. Number of genes used in clustering ranges from top 10 to 100 per cell type ranked by their Cepo statistics. **(B)** For each cell type, correlation of gene statistics calculated from Cepo, Limma, Voom, and EdgeR for each pair of datasets arranged by increasing batch effect as quantified by PVCA.

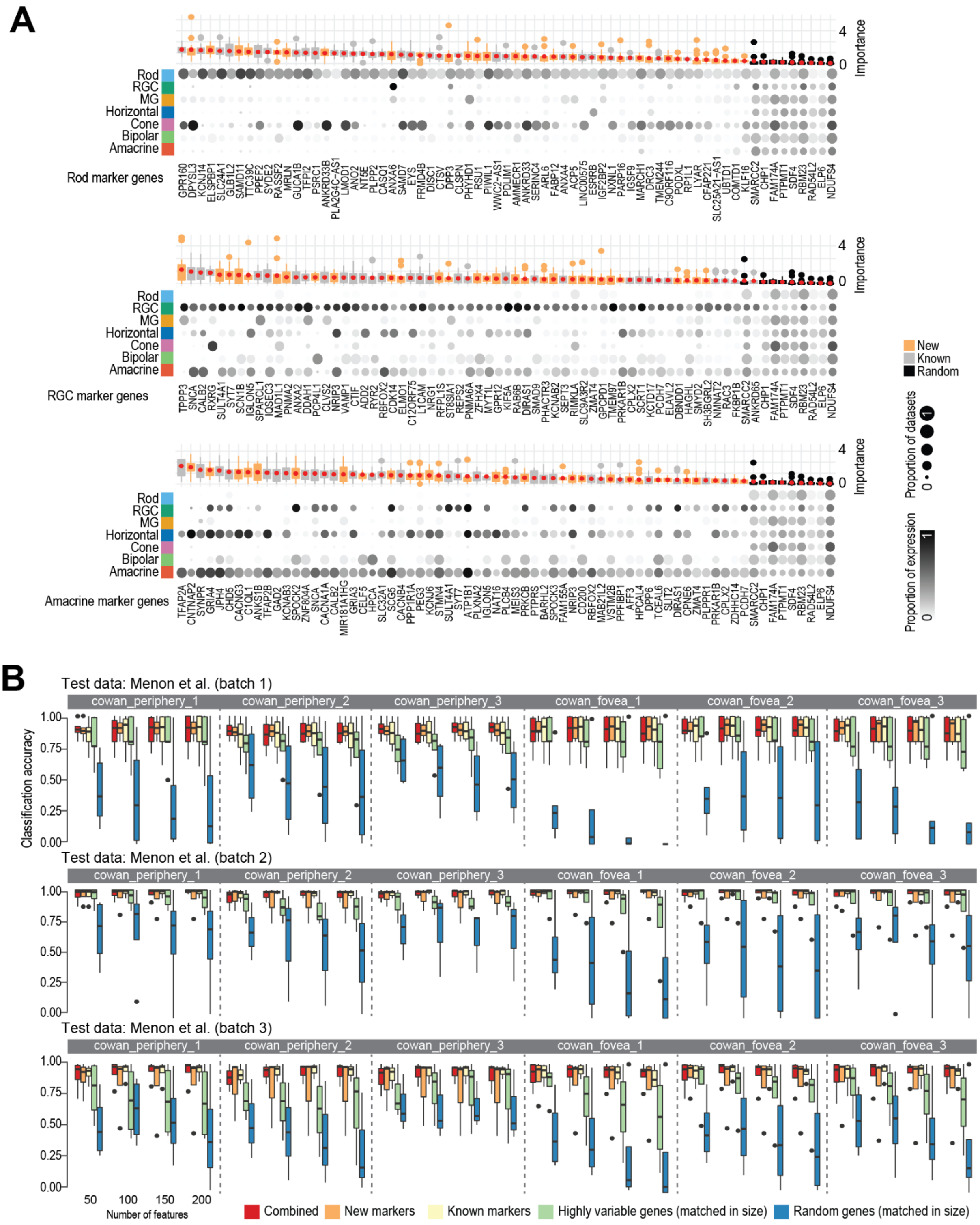


**Figure S3. Cepo identification of cell-type-specific gene markers and their validation on external data. (A)** Cell-type-specific gene markers identified by Cepo for Rods, Ganglion, and Amacrine cells. Proportion of cells expressing each marker in each cell type is represented by the gradient color and the proportion of datasets having each marker expressed is represented by the size of the balloons. Importance scores of gene markers are derived from random forest classification of cells using these markers. Novel markers are highlighted in orange and known markers are in gray. Randomly selected genes (in black) are included as control. **(B)** Classification accuracy of each of the three batches of an independent test data (Menon et al.) from *k*NN classifiers trained on each of the five human retinal datasets and batches using various sets of gene markers (known, new, mixed, highly variable, and random).

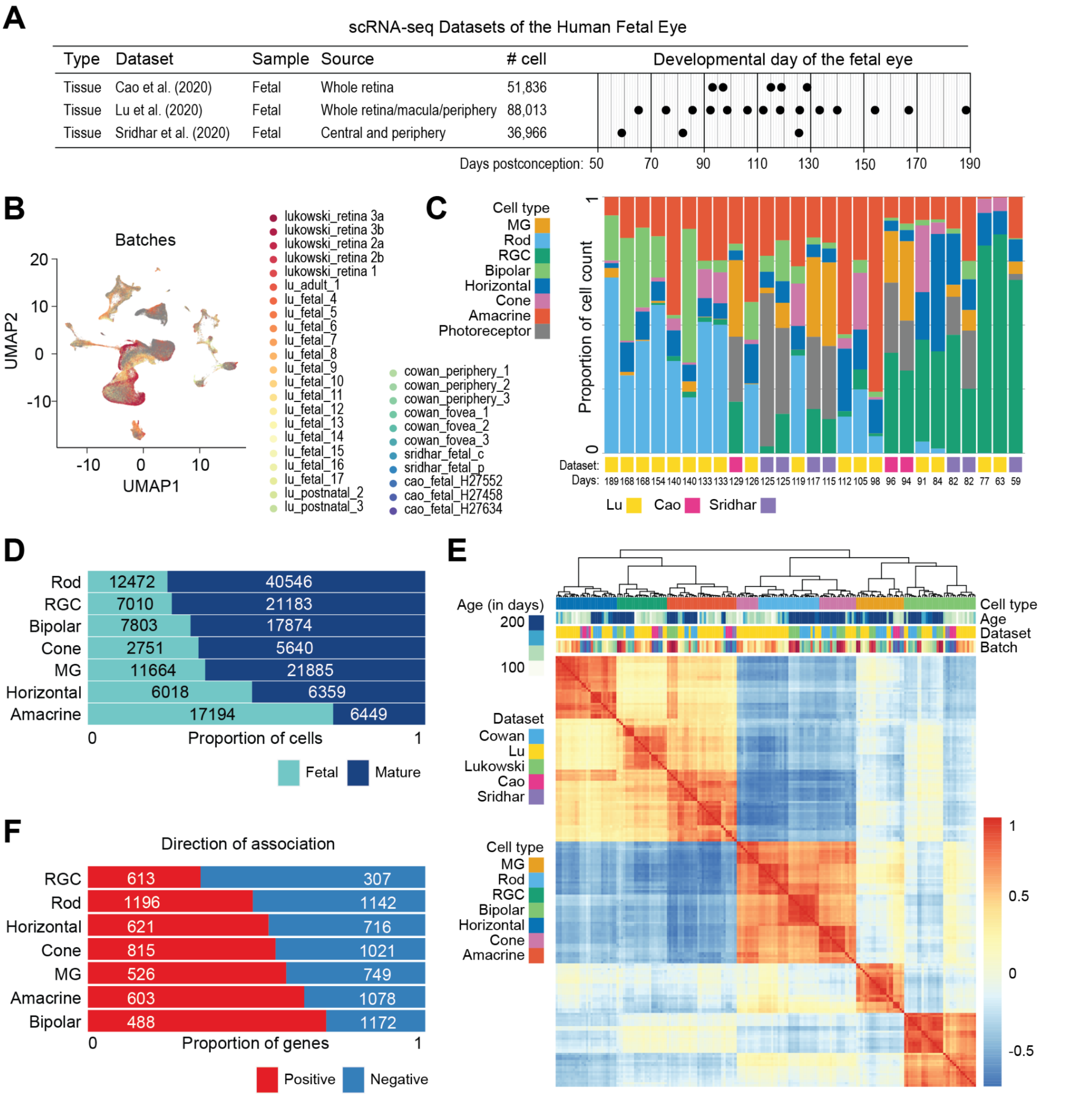


**Figure S4. (A)** Summary of scRNA-seq datasets collected from the retinal tissue. **(B)** UMAP representation of the transcriptomes of single cells from the mature and fetal atlas. Cells are coloured by their batch of origin. **(C)** Proportion of cell types (color coded) in each batch and dataset. Sridhar et al. and Cao et al. contain photoreceptor cells annotated as “Photoreceptors”, which were not resolved into either “Rods” or “Cones. These cells which are not useful in the cross-protocol comparisons were excluded in the study. **(D)** Proportion of mature and fetal cells (color coded) and their total number of cells in each cell type. **(E)** Correlation heatmap of cell-type-specific gene statistics generated from Cepo (Kim et al., 2021) for each cell type across datasets and batches. The heatmap is hierarchically clustered by the similarity of correlation profiles. **(F)** Proportional plot illustrating the direction of association among the significant genes (FDR-adjusted p-value < 0.05). The total number of genes in each set is noted on the right-hand side of the plot.

**A**

scRNA-seq Datasets of Human Retinal Organoids

| Type | Dataset | Cell lines | Age | # cell |
|---|---|---|---|---|
| Organoid | This paper | UCLOOi017-A-1 (017-A-1) and HPSI0314i-hoik_1 (HOIK_1) | 210 days | 34,398 |
| Organoid | Cowan et al. (2020) | 01F49i-N-B7 (F49B7) and iPS(IMR90)-4-DL-01 (IMR90) | 210-266 days | 43,857 |
| Organoid | Lu et al. (2020) | IMR90.4 (IMR90) | 59-205 days | 11,343 |
| Organoid | Kallman et al. (2020) | 1013 | 100-170 days | 14,214 |
| Organoid | Sridhar et al. (2020) | H7 BRN3-td Tomato (H7) | 45-205 days | 10,091 |

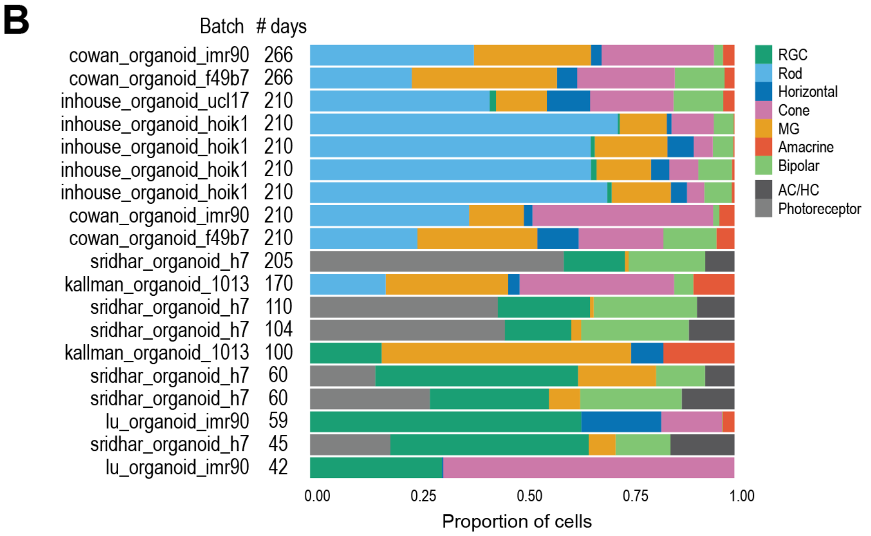


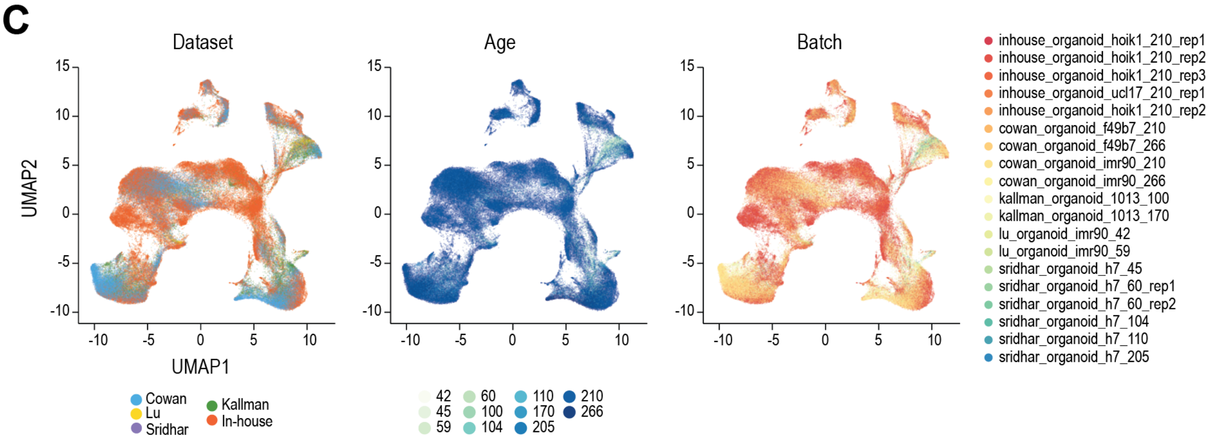


**Figure S5. (A) S**ummary of scRNA-seq datasets collected from the retinal organoid studies. **(B)** Proportion of cell types (color-coded) and total number of cells in each batch and dataset. Sridhar et al. contains photoreceptor cells annotated as “Photoreceptors”, which were not resolved into either “Rods” or “Cones. These cells, which are not useful in the cross-protocol comparisons, were excluded in the study. **(C)** UMAP representation of the transcriptomes of single cells from the organoid atlas. Cells are coloured by the dataset (left), the age until which the organoid was cultured (middle), and their origin of batch (right).

**Table S1**. Summary of public datasets used in this study.

| Dataset | Data source or GEO Accession | Samples | # cells |
|---|---|---|---|
| Cao et al. (2020) | GSE156793 | Fetal | 51,836 |
| Cowan et al. (2020) | https://data.mendeley.com/datasets/sm67hr5bpm/1 | Mature | 51,034 |
| | | Organoid | 43,857 |
| Kallman et al. (2020) | GSE143669 | Organoid | 14,214 |
| Lu et al. (2020) | GSE122970 | Mature | 19,000 |
| | GSE116106 | Fetal | 88,013 |
| | GSE138002 | Organoid | 11,343 |
| Lukowski et al. (2019) | https://zenodo.org/record/5515631#.YUZMWGYzYmI | Mature | 20,009 |
| Sridhar et al. (2020) | GSE142526 | Fetal | 36,966 |
| | | Organoid | 10,091 |

**Table S2**. Comparison of media compositions of human retinal organoid differentiation protocols analysed in this study.

| Reference | Approach | | Basic media composition/Developmental Stage | | | | | | | Isolation/ Dissection |
|---|---|---|---|---|---|---|---|---|---|---|
| | | | hPSC | | Neuroinduction | Retinal Differentiation | | Retinal Maturation | | |
| Gonzalez-Cordero et al. 2017* | Unguided | 2D-3D | E8 | E6 | Advanced DMEM/F12-N2 | DMEM/F12-B27 | DMEM/F12-B27+FBS | Advanced DMEM/F12-B27+FBS +1µM RA | Advanced DMEM/F12-B27-N2 +0.5µM RA *+/- FBS | Day 25-35 |
| West et al. 2022 | Unguided | 2D-3D | E8 | E6 | Advanced DMEM/F12-N2 | DMEM/F12-B27 | DMEM/F12-B27+FBS | Advanced DMEM/F12-B27+FBS +1µM RA | Advanced DMEM/F12-B27+N2 + 7mM glucose + lipids 1:1000 +0.5µM RA | Day 25-35 |
| Cowan et al. 2020 | Unguided | 3D-2D-3D | mTeSR1 | | DMEM/F12-B27 | DMEM/F12-B27 +FBS | | DMEM/F12-B27 +FBS +1µM RA | DMEM/F12-N2 +0.5µM RA | Day 28-32 |
| Sridhar et al. 2020 | Unguided | 3D-2D-3D | mTeSR1/ StemFlex | | DMEM/F12-N2 | DMEM/F12-N2 +FBS | | DMEM/DMEM-F12-B27 +FBS | | Day 18-20 |
| Kallman et al. 2020 | Unguided | 3D-2D-3D | mTeSR1/ StemFlex | | DMEM/F12-N2 | DMEM/F12-B27 | | DMEM/F12-B27 +FBS + lipids +1µM RA | | Day 25-30 |
| Lu et al. 2020 | Guided | 3D | mTeSR1 | | DMEM/E6-B27+1%Matrigel +IWR-1 +SAG | DMEM/F12-B27+FBS +0.5µM RA | | DMEM/F12-B27+FBS +0.5µM RA +10µM DAPT | | Day 10-14 |

hPSC = Human pluripotent stem cell, RA = Retinoic acid, E8 = Essential 8; E6 = Essential 6; * = modifications made to published protocol for this study

**Table S3**. Antibodies used for immunohistochemistry

| Antigen | Host species | Concentration | Supplier |
|---|---|---|---|
| Rhodopsin | mouse | 1 in 1000 | Sigma (04-486) |
| Arrestin 3 | rabbit | 1 in 200 | Abcam (ab189437) |
| Thy1 | rabbit | 1 in 200 | Abcam (ab13350) |
| PKCa | rabbit | 1 in 50 | Thermo (MA1-157) |
| Calretinin | rabbit | 1 in 200 | Abcam (ab702) |
| CRALBP | rat | 1 in 200 | Thermo (MA1-8813) |
| Prox1 | rabbit | 1 in 200 | Merck (ab5475) |
